# A single phospho-LCK flow-cytometry readout predicts dasatinib sensitivity in paediatric T-cell acute lymphoblastic leukaemia

**DOI:** 10.1101/2025.11.04.684929

**Authors:** AA Poll, Y Shi, K Rauwolf, B Bornhauser, A Kulozik, JP Bourquin, JAE Irving, FW van Delft

## Abstract

Children with T-cell acute lymphoblastic leukaemia (T-ALL) who relapse or fail induction have poor outcomes. A subset of cases shows glucocorticoid resistance reversible by dasatinib through inhibition of LCK-dependent signalling. To identify a practical biomarker of dasatinib response, we compared *in-vitro* drug sensitivity with basal phosphorylation of pre-TCR pathway proteins in 28 paediatric T-ALL patient-derived xenografts (PDX). Phospho-flow cytometry quantified LCK (pY394), ZAP70 (pY319) and CD3ζ (pY142), normalised to internal controls. Dasatinib IC_50_ values correlated significantly with pLCK and pZAP70, with pLCK providing the best single-marker performance across clinically relevant thresholds. Logistic, ROC and precision-recall analyses confirmed that pLCK alone achieved excellent classification (AUC ≥ 0.9), while multivariate models added minimal predictive value. Model selection using LASSO and Bayesian Information Criterion further supported pLCK as the dominant predictor. These findings establish pLCK as a robust, scalable biomarker of dasatinib sensitivity suitable for diagnostic integration. A multicentre international validation programme is underway to harmonise pLCK assay protocols, expand testing across biobanks, and assess clinical feasibility in newly diagnosed and relapsed patients within the ALLTogether and HEM-iSMART trials respectively.

## Manuscript

To the Editor,

Children with T-cell acute lymphoblastic leukaemia (T-ALL) who fail induction or relapse have poor outcomes.^1^ Dasatinib can reverse glucocorticoid resistance in a subset of paediatric T-ALL by inhibiting LCK-dependent signalling ^2^, and is being clinically tested (e.g., SJALL23T^3^, HEM-iSMART^4^).

The international HEM-iSMART trial NCT05751044 currently uses a single centre clinically approved and validated *in vitro* co-culture system to assess dasatinib sensitivity in fresh or frozen patient samples^5,6^. The logistics of sending diagnostic samples across Europe means that quantity and quality of samples to be tested might not always be optimal. We hypothesised that integration of an intracellular phosphorylation marker into the routine diagnostic flow cytometry antibody panel could also reliably predict dasatinib sensitivity.

To investigate the correlation between *in vitro* dasatinib sensitivity and phospho-flow cytometry, we profiled a panel of paediatric T-ALL patient-derived xenografts (PDX, n = 28) for basal phosphorylation of proteins in the pre T-Cell Receptor pathway, namely LCK (pY394), ZAP70 (pY319), and CD3ζ (pY142). *Ex vivo* dasatinib sensitivity was quantified as IC_50_ for each PDX sample co-cultured with OP9-DL1 and labelled “sensitive” or “resistant” across a range of thresholds which reflect biological heterogeneity and clinically achievable exposures; 10 nM hypersensitive, 100 nM literature standard^2,7,8^, 1000 nM approximate clinical ceiling^9^. Cells were fixed in 1% formaldehyde, permeabilised using a saponin-based buffer (BD Biosciences PermWash Buffer Cat# 554723), stained with commercially available antibodies against each phospho-target plus CD7 and Thermo LiveDead Violet (Table 1), and analysed on an Attune NxT Flow Cytometer. Events were gated on live human blasts and normalised to the positive control cell line KOPT-K1^10^ and negative Fluorescence Minus One controls^11^ to minimise batch effects.

**Table 1.** Antibodies and dyes. Antibodies used in analytical steps to predict dasatinib sensitivity are listed as Readout. Antibodies and dyes used in cell gating only are listed as Gating.

| TARGET | PHOSPHO-SITE | CLONE | FLUOROPHORE | MANUFACTURER | CAT# | DILUTION |
| --- | --- | --- | --- | --- | --- | --- |
| *Readout* |  |  |  |  |  |  |
| **LCK** | Tyr394 | A18002D | PE | BioLegend | 933104 | 1:20 |
| **ZAP70** | Tyr319 | 1503310 | PE/Cyanine7 | BioLegend | 683708 | 1:20 |
| **CD3ζ** | Tyr142 | K25-407.69 | AlexFluor®647 | BD Biosciences | 558489 | 1:5 |
| *Gating* |  |  |  |  |  |  |
| **CD7** | - | CD7-6B7 | FITC | Biolegend | 343104 | 1:20 |
| **LIVE/DEAD** | - | - | Violet | Invitrogen | L34964 | 1:1800 |

Linear regression showed significant correlation between dasatinib IC_50_ and both pLCK and pZAP70 (p < 0.001, Figure 1A), with pLCK demonstrating the slightly better fit (R^2^ = 0.44 vs 0.4). Univariate logistic regression classifying markers at each threshold (Figure 1B) showed that pLCK was the most informative marker across thresholds, visualised by intuitive confusion matrices (Figure 1C) that match clinical use-cases. Receiver-operating characteristic (ROC, Figure 1D, E) for pLCK alone showed complete segregation using the hypersensitive threshold (AUC = 1) and remained high (≥0.9) at the literature standard of 100 nM threshold. Precision-recall (PR) curves were also robust (Figure 1 F,G). pZAP70 showed similar predictive power, whilst pCD3ζ only demonstrated signal at selected thresholds.

**Figure 1.**
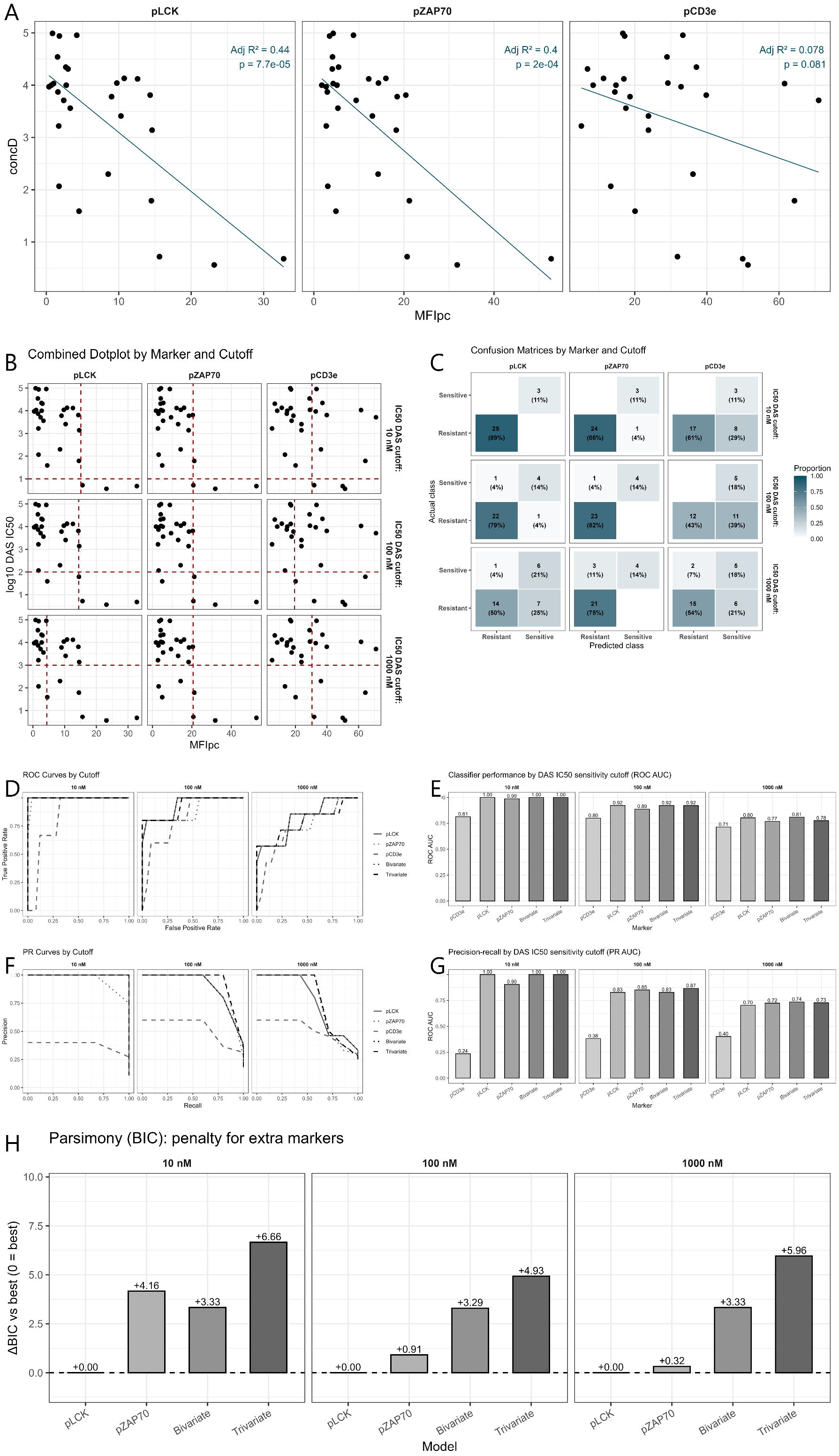
Flow-cytometric predictors of dasatinib sensitivity in paediatric T-ALL PDX. (A) Marker-wise scatter plots (MFIpc vs dasatinib IC50) with fitted slopes. (B) Combined dotplots with IC50 cutoffs (10/100/1,000 nM; red lines) illustrating threshold behaviour. (C) Confusion matrices for single-marker thresholding, showing counts and class proportions. (D) ROC curves by IC50 cutoff comparing pLCK, pZAP70 and pCD3ζ. (E) Bar plots of ROC AUC across markers and cutoffs. (F) Precision–recall curves by IC50 cutoff. (G) Bar plots of PR AUC (same layout). (H) BIC penalty analysis demonstrating that pLCK-only models are preferred over multi-marker models, supporting a parsimonious implementation.

We next examined whether combining markers adds meaningfully to prediction accuracy. Two- and three-marker logistic models produced only marginal gains over pLCK alone, and these were not stable across IC_50_ cutoffs. Least-absolute-shrinkage (LASSO) models^12^ repeatedly selected pLCK as the dominant predictor with little or no contribution from pZAP70 or pCD3ζ in cross-validation. Information-criterion analyses formalised this parsimony: the Bayesian Information Criterion (BIC) ^13^ consistently favoured the pLCK-only model over bivariate or trivariate alternatives (Figure 1H), indicating that any marginal improvements in apparent fit did not overcome the penalty for additional parameters, particularly important when considering clinical implementation.

Taken together, these data support detection of a single-marker, pLCK, as the most reliable and practical response biomarker to identify patients that might benefit from dasatinib treatment — combining strong predictive performance with maximal feasibility for diagnostic laboratories. This approach has several pragmatic advantages, offering:

i. a same-day workflow from sample collection to clinically actionable data;
ii. the possibility to work on fixed samples to reduce handling variability;
iii. wide reagent and infrastructure availability to ensure maximum local implementation, and ;
iv. minimal cell input, with fewer than 100,000 cells required for staining of a separate sample and the possibility to integrate into existing intracellular flow panels.

Guided by these results, we have initiated a multicentre, international, validation programme to validate pLCK as a biomarker for dasatinib sensitivity in paediatric T-ALL. Participating centres have adopted a harmonised SOP (fixation/permeabilisation, antibody clone and titration, gating, normalisation) with centralised QC to ensure cross-site reproducibility. The collaboration will collect pLCK measurements from newly diagnosed patients within the ALLTogether trial^16^ to confirm clinical feasibility and establish a proof of concept dataset to support future implementation.. To confirm diagnostic accuracy, we intend to collect samples from the HEM-iSMART trial patients once open for recruitments in the UK (PI: Dr van Delft) to correlate pLCK as a predictor of dasatinib response with the trial mandated *in vitro* drug response profiling, assessing these against clinical outcomes.

In summary, a minimal phospho-flow assay centred on pLCK stratifies dasatinib response in paediatric T-ALL PDX and is immediately implementable within diagnostic settings. These data provide a rationale and a practical framework for integrating pLCK into ongoing and future trials of dasatinib-containing regimens.

## Acknowledgments

We thank the Newcastle University Flow Cytometry Core Facility, BioImaging Core Facility and Centre for Comparative Biology. The authors acknowledge the contributions of Dr Helen Blair, Samantha Jepson-Gosling and Mankaran Singh in generating patient derived xenograft material for this project. This work was supported by CCLG / Little Princess Trust Project Grant (CCLGA 2022 20 Van Delft), Action Medical Research GN2709 and JGW Patterson Foundation, and makes use of patient-derived xenograft models established at Newcastle University Centre for Comparative Biology under Home Office Project License PPL60/4552 and by researchers with Home Office Personal License under the Animal (Scientific Procedures) Act 1986.

